# Using axenic and gnotobiotic insects to examine the role of different microbes on the development and reproduction of the kissing bug *Rhodnius prolixus* (Hemiptera: Reduviidae)

**DOI:** 10.1101/2022.06.29.497934

**Authors:** Carissa A. Gilliland, Vilas Patel, Ashley C. Dombrowski, Bradley M. Mackett, Kevin J. Vogel

## Abstract

Kissing bugs (Hempitera: Reduviidae) are obligately and exclusively blood feeding insects. Vertebrate blood is thought to provide insufficient B vitamins to insects, which rely on obligate symbiotic relationships with bacteria that provision these nutrients. Kissing bugs harbor environmentally acquired bacteria in their gut lumen, without which they are unable to develop to adulthood. Early experiments identified a single bacterial species, *Rhodococcus rhodnii*, as a symbiont of *Rhodnius prolixus*, but modern studies of the kissing bug microbiome suggest that *R. rhodnii* is not always present or abundant in wild-caught individuals. We asked whether *R. rhodnii* or other bacteria alone could function as symbionts of *R. prolixus*. Bacteria-free (axenic) insects were produced whose microbiome could be experimentally manipulated to produce insects with known microbiomes (gnotobiotic). We found that gnotobiotic insects harboring *R. rhodnii* alone developed faster, had higher survival, and laid more eggs than gnotobiotic *R. prolixus* harboring other bacterial monocultures, including other described symbionts of kissing bugs and several related *Rhodococcus* species. *R. rhodnii* grew to high titer in the guts of *R. prolixus* while other tested species were found at much lower abundance. *Rhodococcus* species tested had nearly identical B vitamin biosynthesis genes, and dietary supplementation of B vitamins had a relatively minor effect on development and survival of gnotobiotic *R. prolixus*. Our results indicate that *R. prolixus* have a higher fitness when harboring *R. rhodnii* than other bacteria tested, and that symbiont B vitamin synthesis is likely a necessary but not sufficient function of gut bacteria in kissing bugs.

## Introduction

All known obligately and exclusively blood feeding arthropods require bacteria to successfully complete development. In most of these cases, the arthropods harbor intracellular symbionts which are passed vertically from mother to offspring with extremely high fidelity, usually via trans-ovarial transmission (Allen et al., 2007; Hosokawa et al., 2010; Pais et al., 2008). Kissing bugs (Hemiptera: Reduviidae: Triatominae) are a group of ∼130 species of insects that obligately and exclusively feed on vertebrate blood, and also require bacteria for successful development (Brecher & Wigglesworth, 1944). In contrast to the symbionts of other exclusively blood feeding arthropods, the symbionts of kissing bugs are not intracellular and are acquired from their environment, often through coprophagy of feces from cohabiting kissing bugs (Brown et al., 2020). Early studies of kissing bug microbiomes suggested that single microbes dominated the microbiome (Brecher & Wigglesworth, 1944; Lake & Friend, 1967; Wigglesworth, 1936). In the model species *Rhodnius prolixus*, an Actinobacteria, *Rhodococcus rhodnii*, was initially identified as the single symbiont inhabiting the anterior midgut, with other kissing bug species harboring distinct microbes(Brecher & Wigglesworth, 1944; Cavanagh & Marsden, 1969; Duncan, 1926; Goodchild, 1955; Gumpert, 1962; Marchette & Hatie, 1965; Weurman, 1946; Wigglesworth, 1936). These studies relied on culture-dependent methods, which we now know underestimated the microbiome.

Explorations of the kissing bug microbiome employing modern 16S amplicon sequencing techniques have shown that the microbiome of most kissing bug species is composed of dozens to hundreds of members (Brown et al., 2020; Carels et al., 2017; da Mota et al., 2012; Díaz-Sánchez et al., 2018; Dumonteil et al., 2018; Gumiel et al., 2015; Kieran et al., 2019; Mann et al., 2020; Montoya-Porras et al., 2018; Oliveira et al., 2018; Rodríguez-Ruano et al., 2018; Waltmann et al., 2019). Differences in species tested, sample size, sampling methodology, and analysis have made comparisons across studies difficult, though several trends are apparent. Studies that have repeatedly sampled the same population and carefully controlled for insect developmental stage have shown that the microbiome shifts through development, becoming progressively less diverse over the life of the insect(Brown et al., 2020; Rodríguez-Ruano et al., 2018). Secondly, Actinobacteria are often found as dominant members of the microbiome (Brown et al., 2020; Carels et al., 2017; Dumonteil et al., 2018; Gumiel et al., 2015; Montoya-Porras et al., 2018; Oliveira et al., 2018). These microbiome surveys suggest that triatomine symbionts lie between the seemingly random acquisition of environmental bacteria as seen in seen in larval mosquitoes (Coon et al., 2016, 2014) and the establishment of specific monoxenic *Burkholderia* symbionts of *Riptoris* sp. (Kikuchi et al., 2007).

In other obligately hematophagous insects, a single species of symbiont is sufficient to support host development. Early studies indicate that *R. rhodnii* is a sufficient symbiont for *R. prolixus* on its own (Brecher & Wigglesworth, 1944; Lake & Friend, 1967), but many of these studies lacked modern tools to assure that their insects were indeed gnotobiotic or axenic. When considering the culture-independent surveys, the lack of a consistent microbiome among individuals within a species suggests that a number of different bacteria can function as symbionts of kissing bugs. Thus, the identity of bacteria that can function as symbionts in this system is unknown.

Another unresolved question in this system is the function of the symbionts. Initial studies implied that the symbionts were necessary for B vitamin synthesis, similar to other obligately hematophagous insects (Baines, 1956; Duncan, 1926). The ability of *R. rhodnii* to synthesize thiamine (B_1_) and folic acid (B_9_) was established by 1960 (Harington, 1960), while the draft genome of *R. rhodnii* identified genes involved in synthesis of B vitamins (Pachebat et al., 2013). However, experimental studies on the role of B vitamins provided mixed evidence for their role in the symbiosis. Lake and Friend (Lake & Friend, 1968) used artificial diets with varying B vitamin composition to show that loss of any B vitamin had a negative effect on molting success, but that even insects reared on a diet containing all the B vitamins did not molt as successfully as bugs harboring symbionts. Hill et al. (1976) developed B vitamin auxotrophic strains of *R. rhodnii* and tested their effect as sole symbionts of *R. prolixus*. They found that for all auxotrophic strains, molting of insects was higher than for aposymbiotic insects. From these data, they concluded that while B vitamin biosynthesis is potentially a necessary feature of a successful symbiont of *R. prolixus*, it may not be sufficient to fully support development of *R. prolixus*.

In this study we focused on members of the genus *Rhodococcus* to assess whether other species in this genus could function as symbionts of *R. prolixus*. These Gram-positive bacteria inhabit diverse environments and ecological niches. Many species in this genus are of interest due to their abilities to break down complex organic compounds including industrial wastes and pollutants (Bell et al., 1999) and have been isolated from diverse environments such as waste water sludge, soil, and aquatic sediments (Garrido-Sanz et al., 2020). One of the most well-studied species of the genus is *R. hoagii*/*equii*, a pathogen of foals, other livestock, and an important pathogen of immunocompromised humans (Mosser & Hondalus, 1996). In addition to *R. rhodnii*, other species within the genus have been identified as symbionts. *R. triatomae* was isolated from the gut of the kissing bug *Triatoma infestans* (Yassin, 2005), while *Rhodococcus* sp. UM008 was isolated from the renal tissue of the skate *Leucoraja ocellata* and may be involved in steroid biosynthesis (Wiens et al., 2016), and *Rhodococcus* WMMA185 was isolated from a sponge, though whether it is strictly host-associated has not been determined (Adnani et al., 2016).

The repeated evolution of symbiosis and host association in this genus led us to ask whether other species could function as symbionts of *R. prolixus*. We examined the effects of 5 *Rhodococcus* species on *R. prolixus* development and reproduction, and compared this to another Gram-positive bacterium (*Micrococcus luteus*) and *Escherichia coli*, which has previously been determined to rescue development of axenic mosquito larvae (Coon et al., 2014). We also examined available genome sequences of *Rhodococcus* sp. to assess whether each tested strain was able to synthesize B vitamins.

## Materials and Methods

### Insect colonies

*Rhodnius prolixus* were obtained from the lab of Dr. Ellen Dotson at the Centers for Disease Control and Prevention through BEI Resources. Insects were housed in an environmental chamber maintained at 28° C with a 12h L:12h D photoperiod and 80% RH. General colony insects were maintained in 1L Nalgene containers with a mesh lid, cardstock folded to allow insects to reach the top of the container, and a 10 cm diameter filter paper on the bottom of the container. Insects were kept in cohorts of several hundred insects per container (1st -3rd instar), ∼100 insects per container (4th-5th instar) or ∼50 insects/container (adults). Colony insects were fed defibrinated rabbit blood (Hemostat Laboratories, Dixon, CA) inoculated with 10^6^ CFU/ml of *R. rhodnii* through an artificial membrane feeder. The feeder consisted of a water-jacketed glass bell with a latex membrane spread over the opening which was attached to a recirculating water bath heated to 37° C. Nymphs were fed bi-weekly for 3 h in the dark at 25° C. Following the blood meal, insects were sorted based on their feeding. Adults were blood fed bi-weekly and eggs were collected following each feed.

### Bacteria

To select bacteria for experiments, 116 *Rhodococcus* genomes classified as having assemblies at either “complete” or “scaffold” level were downloaded from NCBI (table S1). Four genomes from species in the sister genus *Nocardia* were selected as outgroups. Protein sequences from 30 housekeeping genes were extracted from the genomes (table S1) and aligned individually using MAFFT (Katoh et al., 2002) with the –linsi option. Aligned sequences were then concatenated and low-quality regions removed from the concatenated alignment (see supplemental data file S1). The alignment was used to build a phylogeny of the chosen genomes using PhyloBayes-MPI (Lartillot et al., 2009) using the default options (“-cat -gtr”). Two chains were run and used as input to bpcomp with a burn-in of 100 generations and sampling every 10 trees to generate a consensus tree. The consensus tree (figure S1) was visualized with Figtree (https://github.com/rambaut/figtree/releases).

*R. rhodnii* and *R. triatomae* were selected based on previous reports of symbiotic associations with kissing bugs (Brecher & Wigglesworth, 1944; Pachebat et al., 2013; Yassin, 2005). Other species were selected based on their representation of major clades of *Rhodococcus* (figure S1) and availability in public repositories. Cultures of *R. rhodnii* were obtained through the American Type Culture Collection (ATCC), *R. triatomae* DSM-44892 and DSM-44893 were obtained from the Deutsche Sammlung von Mikroorganismen und Zellkulturen (DSMZ), *R. rhodochrous* and *R. erythropolis* and *R. koreensis* were obtained from NITE Biological Resource Center (NBRC), *Escherichia coli* MG1655 and *Micrococcus luteus* were gifts of Eric Stabb and Michael Strand, respectively (table S2). Bacteria were cultured on LB plates or liquid media at 28° C. Growth curves were constructed for each bacteria by inoculating 10 ml of sterile LB with a single bacterial colony and measuring optical density (OD) at 400 nm on a Beckman Coulter DU 640 spectrophotometer until the culture reached stationary phase. At each timepoint, bacteria were serially diluted and plated on LB agar to determine the number of colony forming units (CFU) corresponding to a given OD.

### Generation of axenic and gnotobiotic insects

To generate axenic insects, eggs from the colony were collected 4 d after being laid. In a microbiological safety cabinet, eggs were first immersed in 70% ethanol for 5 minutes in a cell strainer, followed by 3 minutes in 10% povidone-iodine solution, then another 5 minute wash in 70% ethanol, followed by three rinses in sterilized deionized water. Washed eggs were left to dry for at least 1 h, then placed in sterile 550 ml glass mason jars with a 25 mm filter paper at the bottom and covered with mesh fabric. These containers were then placed in sterile 1 L screw-top Nalgene containers with a 2.5 cm hole drilled in the lid which was covered with sterile gas-exchange tape. These containers were returned to the general environmental chamber and allowed to hatch for 5 days. Axenic nymphs were separated in the microbiological safety cabinet with sterile forceps into individual sterile mason jars and starved for 7d prior to feeding. All feedings occurred in a closed microbial safety cabinet that had been UV-irradiated for 10 minutes prior to feeding. Nymphs were fed as described above, except feeding bells were autoclaved prior to use, gamma-irradiated latex was used as a membrane, and all feedings were conducted in the microbial safety cabinet with the sash closed. The axenic status of nymphs was tested for each cohort by sacrificing 3-4 nymphs, extracting total DNA using the methods described below, and performing PCR on the DNA using primers targeting highly conserved regions of the bacterial *16S rRNA* gene (table S3). Only cohorts with no amplification of *16S rDNA* were used for further experiments.

Gnotobiotic insects were generated by adding 10^6^ CFU/ml of log-phase bacterial cultures to sterile blood immediately prior to feeding. Fully engorged insects were then sorted into the wells of a 48-, 24, or 12-well sterile polystyrene cell culture plate for observation. Insect development was monitored daily, and insects were re-fed 2 weeks after all insects had molted or died. Subsequent blood meals were either sterile blood (single inoculation experiments) or inoculated with bacteria as above (repeated inoculation experiments). To examine the role of B vitamin provisioning by symbionts, axenic, 4th instar *R. prolixus* were fed blood meals containing a solution of B vitamins (table S4) along with 10^6^ CFU/ml of bacteria. Insects were then monitored for development to adulthood as before.

### Quantification of gut bacteria

Gnotobiotic or axenic nymphs were sampled at 1 d and 5 d post-blood meal in each instar. Whole insects were washed with ethanol and rinsed with autoclaved water, then ground with a pestle in a microcentrifuge tube. Tissues were resuspended in lysis buffer (10 mM EDTA, 50 mM Tris-Cl pH 7.6, 2% Sarkosyl, 100 U Mutanolysin (Sigma Aldrich)) and incubated at 37° C for 1 h, then heated to 62° C for 10 minutes. Proteinase K (25 μg) and lysozyme (5 mg) were then added to the solution and incubated for 1 h at 37° C. Lysate was then washed twice with phenol:chloroform:isoamyl alcohol (25:24:1) and twice with chloroform. DNA was precipitated from the aqueous phase with 1/10th volume of 5 M sodium acetate and 2 volumes of 100% ethanol. DNA pellets were resuspended in 50 μl of 10 mM Tris pH 7.4 and frozen at -20° C. Total genomic DNA was used as template for qPCR targeting single-copy genes of each bacteria using species-specific primers (table S3). Qiagen Quantifast SYBR green qPCR mix was used in a Qiagen Rotogene qPCR machine with a three-step program: an initial 10 minute denaturation at 95° C, then 30 cycles of 95° C for 20 s, 55° C for 45° s, 68° C for 30s, and a final extension at 68° C for 5 minutes, followed by a melt curve analysis. qPCR products were cloned into the pSCA vector of the Agilent StrataClone PCR cloning kit and used to generate absolute copy number standard curves of the target genes, which were used to quantify the copy number of bacterial genomes in insects.

### Egg laying assays

Single female *R. prolixus* from gnotobiotic colonies were fed 2 weeks post-eclosion on either sterile (single inoculation) or inoculated blood (constant inoculation) as before. Females that fed were isolated in individual wells of sterile 6-well polystyrene plates and provided with sterilized cardstock to deposit eggs. Females were returned to the environmental chamber and allowed to deposit eggs for 10 days, after which egg numbers were counted.

### Genome sequencing, assembly, annotation, and comparisons

Cultures of *R. rhodnii* and *R. triatomae* were grown in LB media to log phase, then genomic DNA was extracted using the method described above. Five μg of genomic DNA was used as input for SMRT library preparation using the Express TPK 2.0 kit and v4 sequencing primer. Multiplexed libraries were run on a PacBio Sequel II system at the Georgia Genomics and Bioinformatics Core. Reads were assembled using CANU (Koren et al., 2017), FLYE (Kolmogorov et al., 2019), and ARROW (Korlach et al., 2017). Assembled contigs were annotated using the Prokka annotation pipeline (Seemann, 2014). *Rhodococcus* species used in the growth assays were selected for the availability of high-quality genome sequences (“complete” according to NCBI Microbial Genome tables). Accessions are listed in table S2. We used KEGG (https://www.genome.jp/kegg/) and MetaCyc (https://metacyc.org/) databases to reconstruct B vitamin synthesis pathways from genomes of microbes tested. Orthologous protein groups (“orthogroups”) were determined using Orthofinder (Emms & Kelly, 2019). We performed gene enrichment analysis using Fisher’s exact test implemented in the Blast2GO package. We first selected orthogroups unique to *R. rhodnii* relative to the other species and compared them to all protein coding genes in *R. rhodnii*. We then performed a second enrichment analysis on the orthogroups that are present in all *Rhodococcus* species tested except *R. rhodnii* using the same approach. In this analysis, genes from *R. triatomae* that had orthologs present in all species tested except *R. rhodnii* were compared for functional enrichment against the complete *R. triatomae* genome.

### Data analysis

Development time and survival were analyzed using a Cox proportional hazards model via the R packages Surveminer and Survival. Due to a large number of zero counts, egg laying data were analyzed with a hurdle model implemented in the package pscl, with Tukey post-hoc tests performed with the package emmeans.

## Results

### Bacteria vary in their symbiotic potential

We first tested whether different bacterial species in the gut of *R. prolixus* could influence the development time and survival of the insects. Instar length varied significantly across development irrespective of the bacterium used for inoculation or inoculation regime, with 5th instars generally taking substantially longer than the 1st - 3rd instars (figure 1, table S5; p = 0.002, Cox proportional hazards). Regardless of treatment, developmental times of *R. prolixus* were similar in the 1st and 3rd instars (figure 1, table S5; p = 0.25, 0.37, Cox proportional hazards) suggesting that symbionts play a minimal role in early development. The length of subsequent instars was highly significantly different among treatments and feeding regimes (p < 0.001, Cox proportional hazards test, table S5). Gnotobiotic insects in the single inoculation group developed slower than insects that were repeatedly inoculated during each blood meal, though this was significant only in the 3rd-5th instars (table S5). Nymphs harboring other *Rhodococcus* species exhibited slower molting than *R. rhodnii* nymphs, especially in the later instars, taking roughly 5 days longer to molt in their final instar regardless of inoculation regime.

**Figure 1:**
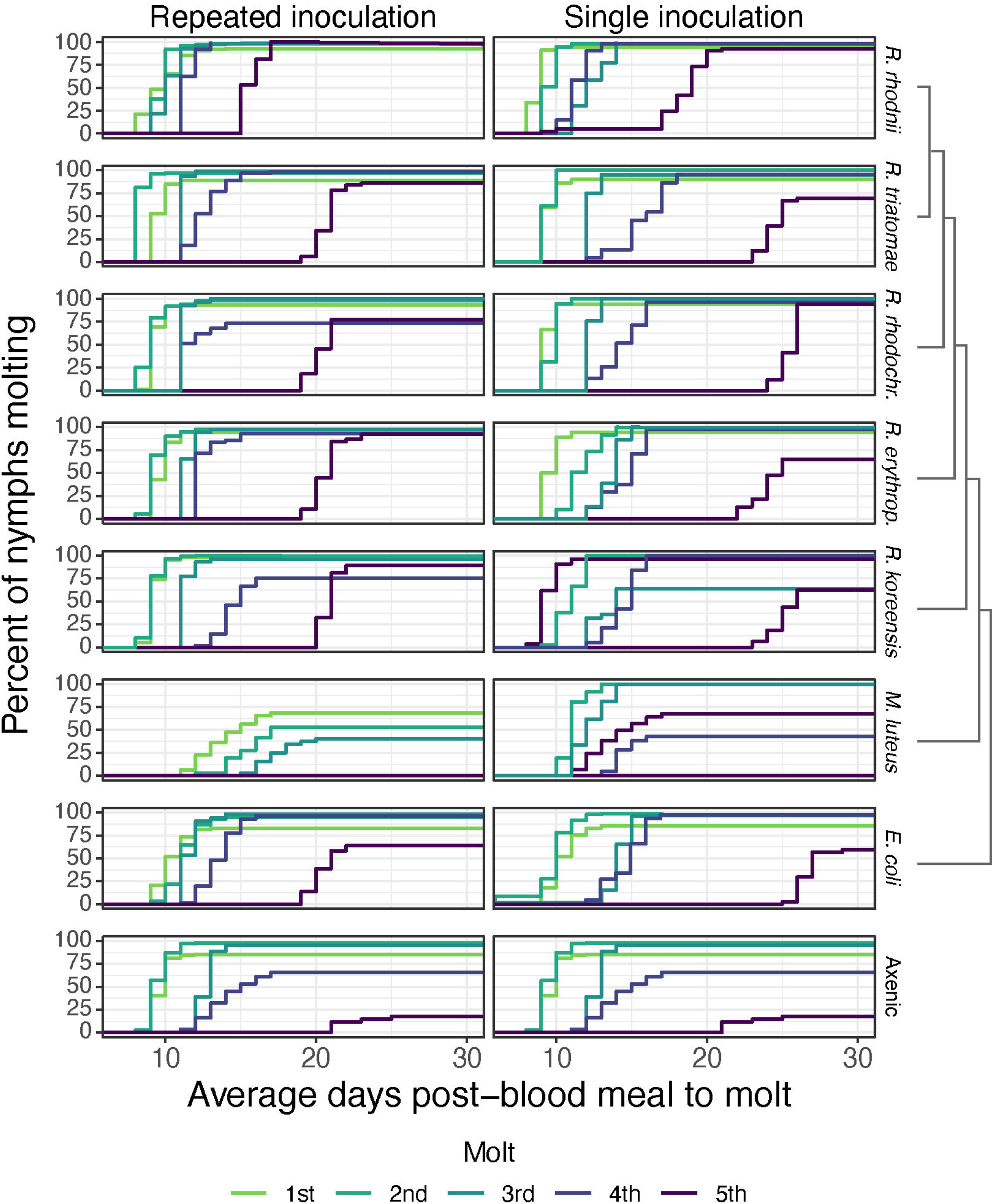
Survival curves of *R. prolixus* inoculated either once (single inoculation) or with each blood meal (repeated inoculation). Insects inoculated with *R. rhodnii* performed significantly better than other treatments (p < 0.05, Cox proportional hazards). Nymphs inoculated with other *Rhodococcus* species and *E. coli* could successfully develop to adulthood, though their development time was longer, and more nymphs failed to molt. Nymphs were blood fed, and only nymphs that successfully fed were retained for the experiment. No nymphs inoculated with *M. luteus* successfully molted to adulthood, and all died prior to the 5th instar. Different dashed lines correspond to different instars. A cladogram next to the bacterial species names indicates relatedness. Nymphs were observed daily to determine the number of days to molt. Nymphs were given 30 days post-blood meal to complete a molt, after which they were classified as having failed to molt.

Survival varied with the bacteria used to generate gnotobiotic *R. prolixus*. As with development time, *R. rhodnii* gnotobiotic nymphs had higher survival regardless of inoculation regime or instar. Survival was high for all *Rhodococcus* sp. and *E. coli* treatments throughout early instars but declined in the 4^th^ and 5^th^ instars. All the *Rhodococcus* gnotobiotics had high survival (> 75%) as 4^th^ and 5^th^ instars in the repeated inoculation regime, though only the *R. erythropolis* gnotobiotics had high survival as 5^th^ instars in the single inoculation regime. The nymphs inoculated with *E. coli* had lower survival as 5^th^ instars regardless of inoculation regime. As expected, axenic insects nearly all died prior to the final molt. Insects inoculated with *M. luteus*, regardless of inoculation regime, died prior to reaching adulthood, suggesting that it cannot function as a symbiont of *R. prolixus*. Together, these data suggest that *R. rhodnii* is a superior symbiont to other *Rhodococcus* species, but that the other species tested, along with *E. coli*, can support development of *R. prolixus* nymphs to adulthood, though at a substantial cost if the bug is not repeatedly inoculated with the bacteria. In contrast, *M. luteus* does not seem capable of functioning as a symbiont.

### R. rhodnii *has consistently high titer*

We hypothesized that symbiotic potential may be correlated with bacterial titer in the gut. To assess this, we generated cohorts of gnotobiotic or axenic nymphs similar to the nymphs used in the developmental experiments. We fed axenic 1st instar nymphs with blood inoculated with 10^6^ CFU/ml while all remaining blood meals were sterile. For each instar, five nymphs from each cohort were sacrificed 1 d and 5 d post-blood meal and used for total DNA extraction. We performed qPCR on the extracted DNA to measure the copy number of a single-copy bacterial gene using primers unique to each bacterial species tested.

Bacterial abundance varied greatly among the species tested and developmental stage (figure 2). *R. rhodnii* titer was consistently higher than other species, quickly reaching ∼10^6^ copies and remaining high throughout development (10^4^ - 10^7^ copies/insect). Titer decreased after each molt, presumably as many bacteria are shed with the contents of the gut prior to molting, then recovering towards the end of each instar. There was a significant decrease in *R. rhodnii* titer between the 2nd and 3rd instar, though its abundance recovered throughout the remaining instars.

**Figure 2:**
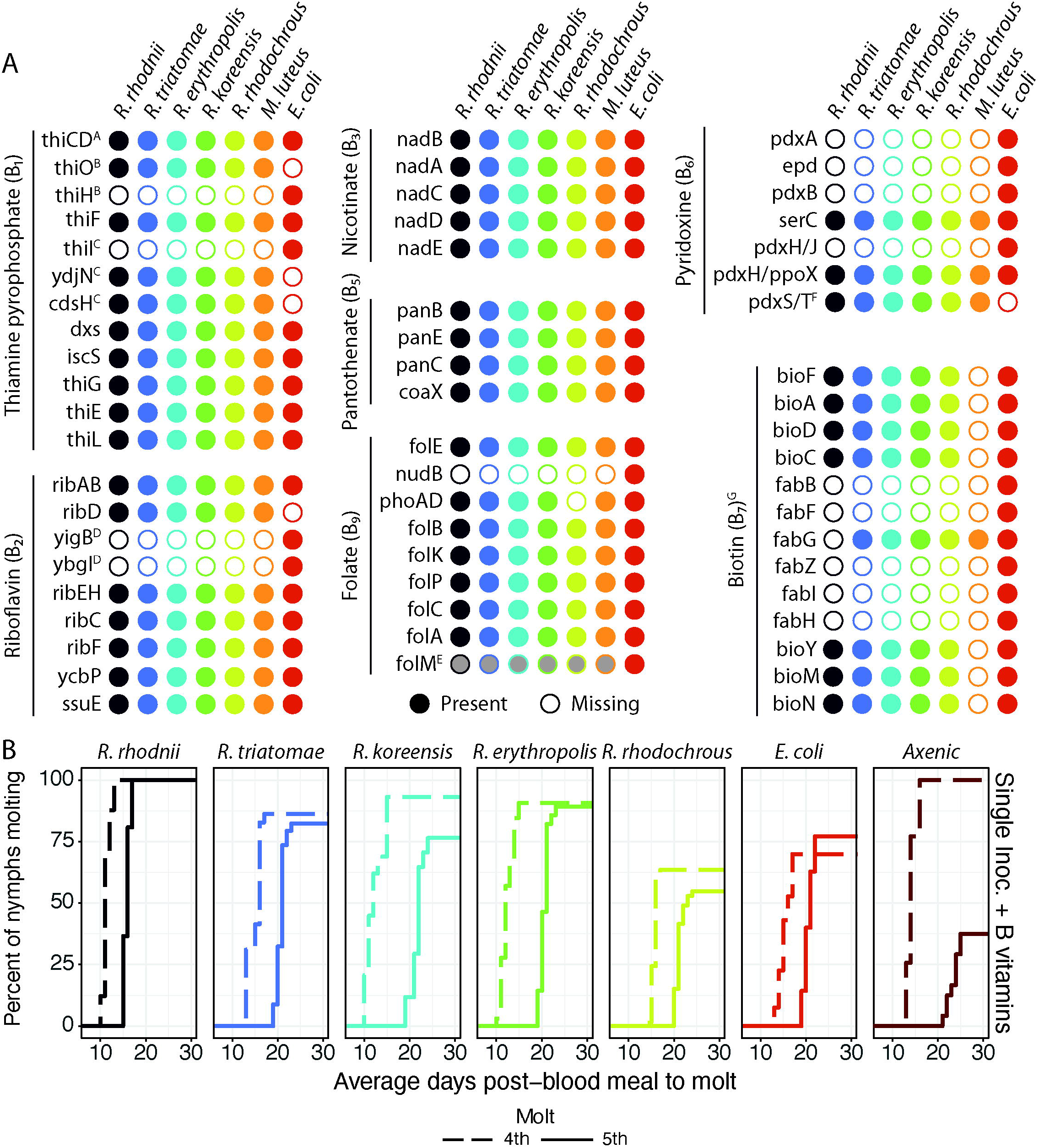
Titer of bacteria in insects from the single inoculation experiments. *R. rhodnii* was found at higher titer than other bacteria at every time point. Bacterial abundance fluctuated over time, decreasing after a molt and subsequently recovering by 5 days PBM. Titer of most bacteria declined after the 2nd instar and failed to recover, though *R. erythropolis* titer remained low but stable throughout development. No insects inoculated with *M. luteus* successfully molted beyond the 4th instar. Bacterial titer was measured using qPCR against the single-copy *gyrB* gene using species-specific primers (table S3). Bacteria were fed at 10^6^ CFU/ml in the first blood meal and all subsequent blood meals were sterile. Titer was measured at 1 and 5 days post blood meal (PBM) by homogenizing whole insects (1st - 3rd instars) or guts only (4th and 5th instars). Shaded area represents 95% confidence interval.

Other bacteria displayed different patterns of abundance. *R. triatomae, R. rhodochrous*, and *R. koreensis* had steady titer for the first two instars but declined rapidly following the 2nd to 3rd instar molt and never recovered (figure 2). The decrease in titer roughly corresponded to lower molting success and delayed development of these gnotobiotic groups of *R. prolixus* (figure 1). The titer of *E. coli* in gnotobiotic *R. prolixus* peaked during the 1st instar then dropped to a low level around 10^2^ copies/insect and remained low throughout the rest of insect development. Similarly, titer of *R. erythropolis* remained near 10^2^ copies/insect, yet a high proportion of these gnotobiotic insects molt to adulthood, whereas other gnotobiotic insects in the single inoculation regime experiments had low success reaching adulthood (figure 1). The titer of *M. luteus* dropped precipitously after inoculation and remained at very low levels until all insects ultimately died. These data suggest that the low survival of *R. prolixus* inoculated with *M. luteus* is unlikely to be due to proliferation or pathogenicity of the bacteria to the insect. Taken together with the developmental data, the effects of different bacteria on development time correlates with bacterial abundance, but the proportion of nymphs successfully molting to adulthood did not always correspond to bacterial titer.

### Gut bacteria influence egg laying

Both single and constant inoculation gnotobiotic *R. prolixus* were assayed for egg production. Following the final molt to adulthood, male and female nymphs were allowed to mate for 2 weeks, then fed a sterile blood meal. Females that fed were separated into wells of a sterile 6-well cell culture plate and provided with autoclaved cardstock to deposit eggs for 10 days. Egg laying was analyzed with a two-step hurdle general linearized model to account for the large number of insects that laid no eggs across treatments. There was a highly significant effect of bacterial treatment (p < 0.001 for all treatments, figure 3, table S6) and for the inoculation regime (p = 0.0054, figure 3, table S6). Insects harboring *R. rhodnii* laid the most eggs regardless of inoculation regime, with those inoculated with *R. erythropolis* laying the next most eggs. The *R. prolixus* inoculated with *E. coli* or repeatedly inoculated with *R. rhodochrous* laid the fewest eggs. *M. luteus* gnotobiotic nymphs did not successfully molt to adulthood, so no egg laying data were available.

**Figure 3:**
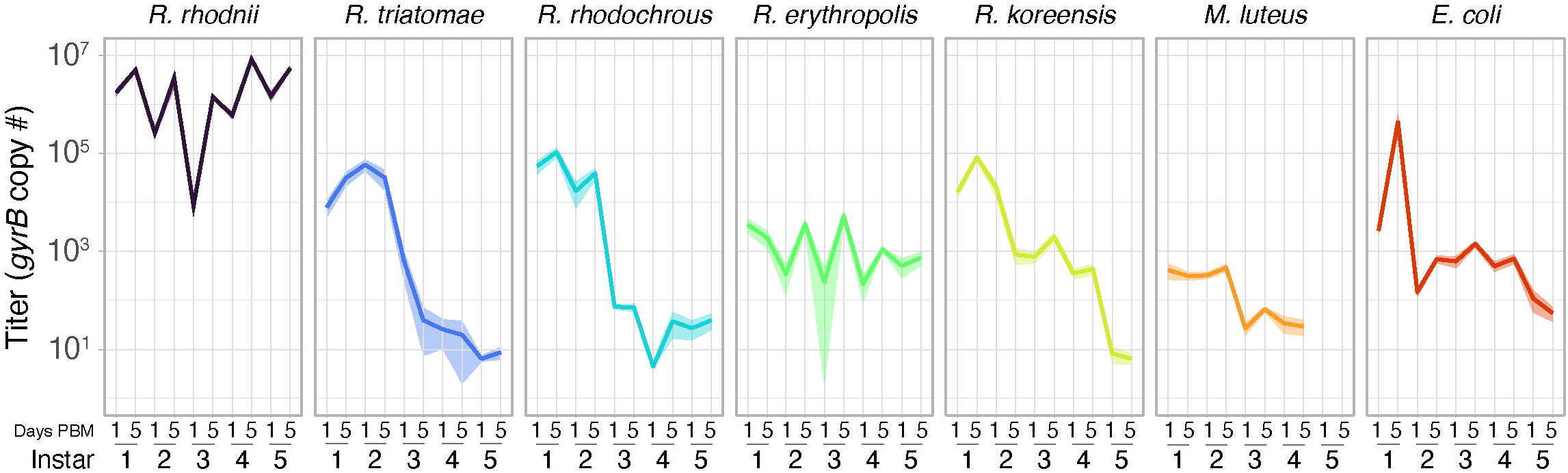
Egg laying of adult female *R. prolixus* by bacterial inoculation. Insects harboring *R. rhodnii* laid more eggs than all treatments except those harboring *R. erythropolis*. Females were blood fed following their final molt and allowed to lay eggs for 10 days post-blood meal. Significant differences were determined using a two-step hurdle generalized linear model followed by a Tukey’s post-hoc test. Treatments not connected by the same letter are significantly different (p< 0.05).

### Genomic comparisons between symbiotic and non-symbiotic Rhodococcus species

The bacteria tested varied in their ability to fulfill a symbiotic role in *R. prolixus. R. rhodnii*-inoculated nymphs performed better in every metric tested (survival, developmental time, and egg production), while *M. luteus* gnotobiotic insects all died prior to adulthood. However, other strains of *Rhodococcus* had mixed success in their ability to function as symbionts of *R. prolixus*, and this did not appear to reflect the phylogenetic distance between bacterial species. To identify potential genes associated with successful symbionts, we explored the genome sequences of the *Rhodococcus* species used in the study. We sequenced the genomes of *R. rhodnii* and *R. triatomae* using the PacBio Sequel system and assembled reads with CANU (Koren et al., 2017) and Flye (Kolmogorov et al., 2019). We successfully produced near-chromosome level assemblies, producing 4 contigs in our assembly of *R. rhodnii* and 1 contig each for two *R. triatomae* strains. RefSeq annotations were produced from these assemblies by the NCBI prokaryotic genome annotation pipeline (PGAP). Assemblies and annotations can be found under the NCBI accessions PRJNA483464 (*R. rhodnii*) and PRJNA606336 (*R. triatomae*).

We first compared the presence of orthologs among the bacteria using OrthoFinder (Emms & Kelly, 2019). Protein coding genes were assigned to phylogenetically determined clusters of orthologs among the genomes, referred to subsequently as orthogroups. Using these data, we assessed the presence or absence of orthogroups between different groups of species. We found a core genome of 2231 orthogroups shared among all species. The genome of *R. rhodnii* encoded 304 unique proteins not found in the other genomes examined, while there were 316 proteins found in the non-*R. rhodnii* species but absent from *R. rhodnii*. We examined functional enrichment of these unique gene sets using Blast2GO and found two clusters that were overrepresented in the unique *R. rhodnii* gene set. In the genes unique to *R. rhodnii*, two functional categories were overrepresented: IPR010982 (12 genes, 4.4% of *R. rhodnii* unique sequences versus 0.55% of genomic sequences, p = 1.87 × 10^−6^, Fisher’s exact test) and IPR001387 (11 genes, 3.8% of *R. rhodnii* unique sequences versus 0.47% of genomic sequences, p = 3.15 × 10^−6^, Fisher’s exact test). IPR001387 and IPRO010982 are hallmarks of the two components of the Cro/C1 bacteriophage repression system that has been shown to suppress transition of bacteriophage λ from the lysogenic to lytic pathways (Schubert et al., 2007). These results prompted us to examine the genome of *R. rhodnii* for phage using the PHASTER server (Arndt et al., 2016; Zhou et al., 2011), which identified 3 complete phage genomes present in the bacterium. Of the other *Rhodococcus* species, only *R. erythropolis* had a complete phage. *R. triatomae, R. koreensis*, and *R. rhodochrous* had none. No functional categories were enriched among the genes present in all non-*R. rhodnii Rhodococcus* genomes but missing from *R. rhodnii*.

We also examined the presence of B vitamin biosynthesis genes among the different species tested in our bioassays (Figure 4A, table S7). While several authors have provided evidence that *R. rhodnii* encodes some genes necessary for B vitamin biosynthesis (Pachebat et al., 2013; Tobias et al., 2020), to date, the presence or absence of complete pathways in the genome has not been examined. All tested *Rhodococcus* strains and *E. coli* encode the complete suite of genes necessary for the *de novo* synthesis of the B vitamins thiamine, riboflavin, nicotinate, pantothenate, pyridoxine, and folate, though in some cases the synthesis pathways differ between the *Rhodococcus* species and *E. coli*. This is most obvious in pyridoxine (B_6_) synthesis, where the genes *pdxABHJ* and *epd* are missing from the *Rhodococcus* species and *M. luteus*. In these species, pyridoxine is synthesized via proteins encoded by *pdxST. R. koreensis* lacks an ortholog to the alkaline phosphatase *phoA/B* involved in folate biosynthesis but encodes numerous other alkaline phosphatases that may perform this reaction. The *Rhodococcus* genomes encode several highly conserved genes for the biosynthesis of biotin via the 8-amino-7-oxononanoate I pathway (*bioFADBC*) but lack the genes necessary to synthesize the pimeloyl coenzyme A moiety, suggesting these bacteria cannot synthesize biotin *de novo*. In contrast, *M. luteus* lacks nearly all genes necessary to synthesize biotin (*bioFADC*). All the *Rhodococcus* species and *M. luteus* encode *bioYNM* which encode a biotin transporter, suggesting that the bacteria can acquire biotin from their environment. These results further suggest that biotin is likely only partially synthesized by these bacteria, relying on external pimeloyl-CoA for *de novo* synthesis or importation of biotin via *bioYNM*. We did not fully consider biosynthesis of cobamides (B12), as these are thought to not be essential to insect development (Dadd, 1973). The invariable presence of the complete synthesis pathways for thiamine, riboflavin, nicotinate, pantothenate, pyridoxine and folate in the *Rhodococcus* species tested suggests that while provisioning one or more of these B vitamins is likely a necessary feature of a successful symbiont of *R. prolixus*, other factors likely underly the variation observed in development, survival, and reproduction among the various gnotobiotic treatments.

**Figure 4:**
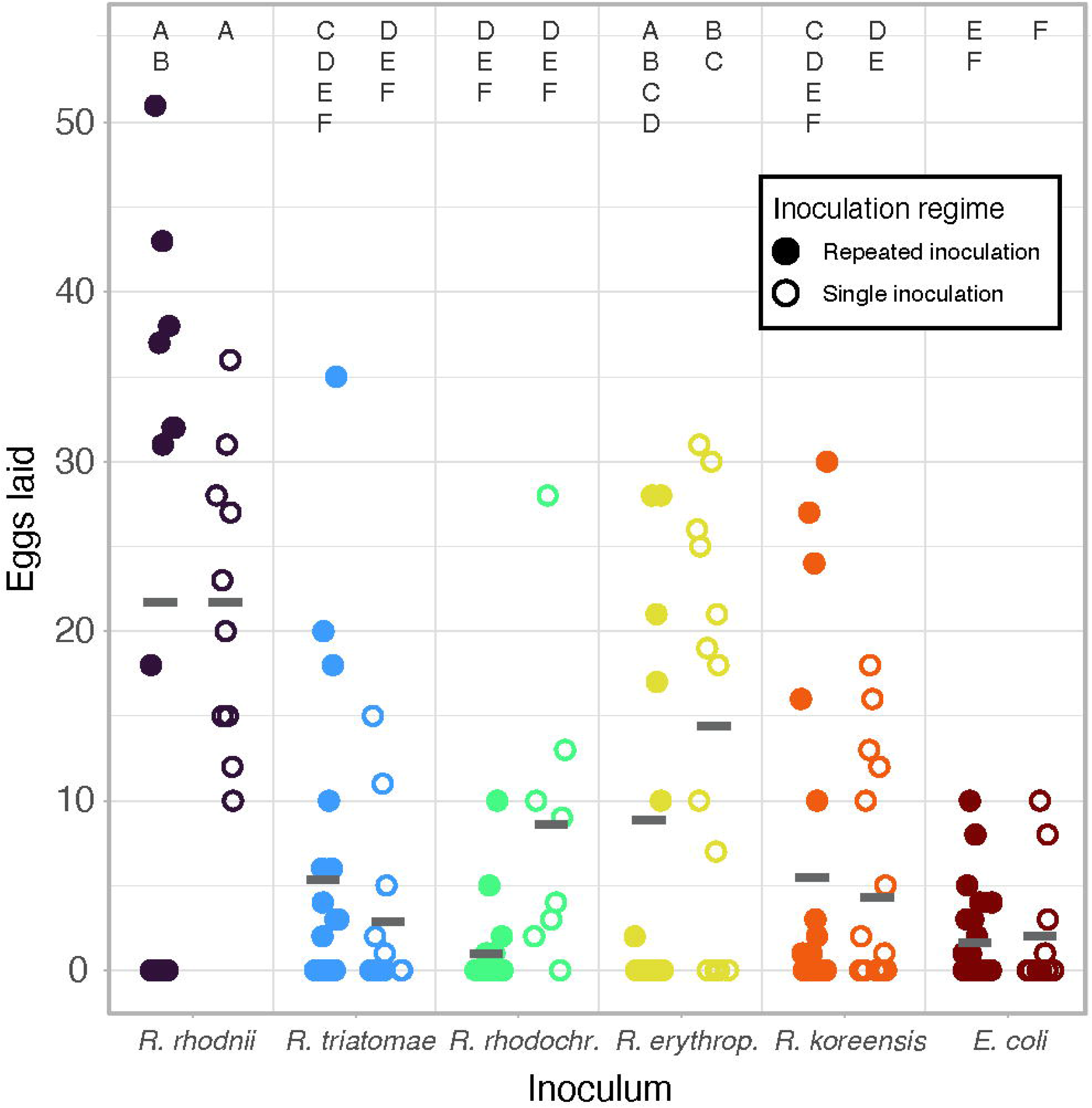
B vitamin biosynthesis potential and effects of B vitamin supplementation on the development of gnotobiotic and axenic *R. prolixus*. A) Presence and absence of core B vitamin biosynthesis genes in the genomes tested. Orthologs were identified through BLAST searchers of genomes and by querying the KEGG and MetaCyc databases. Open circles indicate no ortholog of the enzyme was found in the annotated proteins of the genome. ^A^ *thiCD* is a single bifunctional enzyme in *Rhodococcus* species and *M. luteus*, but is encoded by two separate enzymes in *E. coli*. ^B^ *thiO* is necessary for B1 synthesis using glycine as a precursor while *thiH* is required when tyrosine is the precursor. ^C^ *Rhodococcus sp*. encode *thiI*, but the gene lacks a rhodanase domain which is necessary for sulfurtransferase activity (Martinez-Gomez et al., 2011). In *Salmonenlla enterica*, YdjN and CdsH have been shown to fulfill the sulfurtransferase role of ThiI (Palmer et al., 2014). ^D^ *yigB* and *ybjI* are not required for riboflavin synthesis in *Rhodococcus* or *M. luteus* but are essential for *E. coli* riboflavin biosynthesis. ^E^ FolM is likely not essential for folate synthesis, but rather involved in the production or reduction of 7-8-dihydropterins (de Crécy-Lagard et al., 2007). ^F^ Pyridoxine is synthesized via different pathways in *E. coli* relative to the *Rhodococcus* species and *M. luteus*. In the non-*E. coli* species, pyridoxal 5’-phospate synthase (PdxST) synthesizes B pyridoxal 5’-phosphate from glutamine, D-ribose 5’-phosphate, and glyceraldehyde 3-phosphate. ^G^ While all the species tested except *M. luteus* can synthesize biotin, only *E. coli* can synthesize the pimeloyl-ACP. B) Development of 4^th^ instar gnotobiotic and axenic *R. prolixus* nymphs supplemented with B vitamins in their blood meal.

While our genomic data indicate that the tested *Rhodococcus* species cannot make biotin *de novo*, we directly tested whether the bacteria could be reared without any B vitamins in the media. We found that all species could form colonies within 6 days of inoculation on minimal media plates. To verify that there was no carryover of B vitamins from previous cultures, we picked individual colonies of each species from our initial minimal media plates and streaked them on fresh minimal media plates. Again, we detected growth of all the *Rhodococcus* species tested, suggesting that they are capable of synthesizing biotin without the canonical genes *bioFADBC*.

We then tested whether supplementation with B vitamins via a blood meal could rescue *R. prolixus* development. Axenic or gnotobiotic 4th instar nymphs were provided with a blood meal containing all standard B vitamins (table S4), along with 10^6^ CFU/ml of each bacteria tested. Axenic insects given blood meals supplemented with the B vitamins all successfully molted to 5th instars, while nearly 40% of axenic 5th instars successfully molted to adulthood (Figure 4B), suggesting that B vitamins are partially responsible for the observed failures of axenic nymphs to reach adulthood. The effect of B vitamin supplementation on the growth and molting success of gnotobiotic nymphs was limited, with only slight increases in molting success or reductions in development time. Together with the data from axenic nymphs, these data suggest that B vitamins provisioning by gut bacteria is likely necessary but not sufficient to support optimal insect growth and development.

## Discussion

Initial studies of the kissing bug microbiome implemented culture-dependent manipulations of kissing bugs and specific microbiota (Gumpert, 1962; Lake & Friend, 1968, 1967), but lacked molecular tools to verify axenic or gnotobiotic states, which may explain their conflicting results (Nyirady, 1973). More recent investigations of the kissing bug symbionts have utilized a culture-independent 16S sequencing approach and have focused on characterizing the microbial community of various kissing bug species (Brown et al., 2020; Carels et al., 2017; da Mota et al., 2012; Díaz-Sánchez et al., 2018; Dumonteil et al., 2018; Gumiel et al., 2015; Kieran et al., 2019; Mann et al., 2020; Montoya-Porras et al., 2018; Oliveira et al., 2018; Rodríguez-Ruano et al., 2018; Waltmann et al., 2019). While these studies have broadened our understanding of kissing bug microbiomes, comparisons between studies have been difficult due to different sampling techniques, species used, use of lab versus wild-caught insects, and other confounding variables. These studies are by design, descriptive, and provide limited inference as to which members are symbiotic or csommensal, and what roles specific species play in the biology of the host. As a result, our understanding of the functional roles of microbes in the kissing bug gut remain poorly understood.

We produced bacteria-free, axenic *R. prolixus* via egg surface sterilization and verified their axenic state using culture-independent methods, then generated gnotobiotic insects by providing bacterial inocula through a blood meal. Our methods allowed a high degree of control over the kissing bug microbiome unlike earlier experiments as well as a platform to conduct experiments on the roles of specific microbes in kissing bugs. Consistent with previous studies, axenic insects had high mortality and rarely reached adulthood. *R. rhodnii*-gnotobiotic *R. prolixus* developed faster and with higher success than other gnotobiotic insects. Gnotobiotic insects inoculated with other *Rhodococcus* species and *E. coli* had increased developmental times and decreased survival, which was particularly pronounced in the final nymphal instar. However, many of the insects from these gnotobiotic colonies did eventually reach adulthood, though most laid significantly fewer eggs than insects harboring *R. rhodnii*. In contrast, *M. luteus* proved to be a poor symbiont with no gnotobiotic insects surviving beyond the 4th instar. While *R. rhodnii* is not consistently recovered from 16S surveys of triatomine microbiomes, our data suggest that it confers a significant fitness advantage to *R. prolixus*. We are currently examining whether the fitness benefits or *R. rhodnii* to *R. prolixus* persist when additional microbes are present in the gut, and whether this fitness benefit extends to other species of kissing bug.

Culture-independent 16S surveys of kissing bug microbiomes suggest that nymphs acquire bacteria from their environment which then persist in the insect midgut throughout development (Brown et al., 2020; Rodríguez-Ruano et al., 2018). The ability of the bacterial species tested to persist in the kissing bug gut may reflect the fitness differences observed between gnotobiotic insects harboring different bacteria. We found that many of the bacteria tested persist in the gut throughout development, but at significantly lower levels than *R. rhodnii*. Repeated inoculation of insects with a high titer of bacteria in every blood meal did result in increased survival and faster development, but ultimately did not rescue egg production. We suspected that lower titer of bacteria may have led to lower amounts of B vitamin provisioning by symbionts, and we found that supplementation of blood meals with B vitamins did increase survival and shorten developmental times, but not to the same degree that harboring *R. rhodnii* without supplementation did.

Given that *R. triatomae* it was isolated from another kissing bug species and that it was the closest known relative of *R. rhodnii*, we expected *R. triatomae-*inoculated insects to have a fitness most like those inoculated with *R. rhodnii*, but gnotobiotic insects inoculated with *R. triatomae* were more similar to those harboring other *Rhodococcus* species and not to *R. rhodnii*-inoculated insects. *R. rhodnii* and *R. triatomae* have identical B vitamin synthesis pathways and share nearly 80% of their orthogroups. One possible explanation is the dramatic decrease in *R. triatomae* titer in *R. prolixus*, as fewer than 100 copies of *R. triatomae* were recovered from late-instar insects compared with > 10^6^ copies of *R. rhodnii*. We are currently investigating whether this is due to an inability of the bacteria to survive in the gut environment or factors such as immune activation by *R. triatomae* in *R. prolixus*.

We identified two gene clusters that were over-represented in the unique genes of *R. rhodnii*, and both were associated with regulation of the lytic-lysogenic cycle of bacteriophages. This led us to examine the presence of phage in the *Rhodococcus* genomes tested, and we discovered that *R. rhodnii* had three intact phage integrated into the genome. In contrast, *R. triatomae, R. koreensis*, and *R. rhodochrous* did not encode any complete phage while *R. erythropolis* shared the *Rhodococcus* “sleepyhead” phage (NC_048782.1) with *R. rhodnii*. Whether the phages of *R. rhodnii* contribute to its symbiotic abilities in *R. prolixus* is unknown, but in other symbiotic systems phage play a central role in symbiotic function. The phages of *Hamiltonella defensa*, a secondary symbiont of aphids, encode toxins that likely explain variation in the ability of different strains of the bacteria to kill attacking parasitoid wasps(Boyd et al., 2021; Brandt et al., 2017; Lynn-Bell et al., 2019; Oliver et al., 2009).

Historically, the role of bacterial symbionts in the biology of kissing bugs has been debated, with some researchers suggesting that the necessity of symbionts for kissing bug development was not directly related to B vitamin provisioning (Hill et al., 1976), while others argued that nutrient provisioning was the primary function of symbionts in this system (Lake & Friend, 1968). Our results indicate that B vitamin provisioning is likely necessary when *R. prolixus* is fed on rabbits, though we did not test other sources of blood. However, there was substantial variation in symbiotic potential among the other bacteria tested, despite an apparently uniform ability to synthesize B vitamins. Likewise, addition of B vitamins to blood meals of axenic insects did not fully rescue their development, and *M. luteus*, which cannot synthesize biotin, failed to support *R. prolixus* development. From these experiments we conclude that B vitamin provisioning is likely a necessary function of symbiotic bacteria in *R. prolixus* but is not alone sufficient. Studies using metagenomics approaches in *R. prolixus* with unmanipulated microbiomes suggest that a number of bacterial species may cooperate to provide the host with B vitamins (Tobias et al., 2020). Tobias and colleagues found that among the reads sequenced from a lab colony of *R. prolixus*, many of the B vitamin biosynthesis genes mapped to a species belonging to the genus *Dickeya*, suggesting that in their colony, *R. rhodnii* was not the only bacterium synthesizing B vitamins for the host. Their results and the variable presence of *R. rhodnii* in field-collected *R. prolixus* hint at potential cooperation or perhaps competition between symbionts in the *R. prolixus* gut.

The role of symbiotic bacteria in kissing bugs is likely complex and multifaceted, with symbionts fulfilling nutritional as well as other roles in host physiology. Here we attempted to identify individual bacterial strains that are capable of functioning as symbionts and determined that *R. rhodnii* was a superior symbiont to other species tested, despite conservation of B vitamin biosynthesis genes among tested species. Further studies of the interactions between *R. rhodnii* and other bacteria will illuminate these dynamics, providing additional insights into their function in the host.

## Acknowledgements

The authors would like to thank Logan Harrell and Betsy Jackson for their assistance rearing insects. Alice Sutcliffe provided invaluable advice on establishing and maintaining colonies of *R. prolixus*. We also thank Dr. Ellen Dotson of the Centers for Disease Control and Prevention for her gift of the insects to start the *R. prolixus* colony. This research was supported by startup funds from the University of Georgia College of Agricultural and Environmental Sciences to KJV.

## Supplemental Material

Tables:

S1: Accession numbers of genomes and genes for phylogeny

S2: Accessions of bacterial strains used

S3: Primers used for quantification of bacteria

S4: B vitamin concentrations for feeding experiments

S5: Cox PH statistics

S6: Hurdle model statistics

S7: Accessions of B vitamin biosynthesis genes

Figures/data:

Data S1: Alignment of genes for phylogeny

Figure S1: Phylogeny of *Rhodococcus* species

## Author Contributions

CAG, VP, and KJV designed the research, CAG, VP, ACD, BM, and KJV performed the research. CAG, VP, and KJV analyzed the data. KJV wrote the paper.

