## Supplementary figures and images for "Using axenic and gnotobiotic insects to examine the role of different microbes on the development and reproduction of the kissing bug *Rhodnius prolixus* (Hemiptera: Reduviidae)"

### Supplemental figure s1

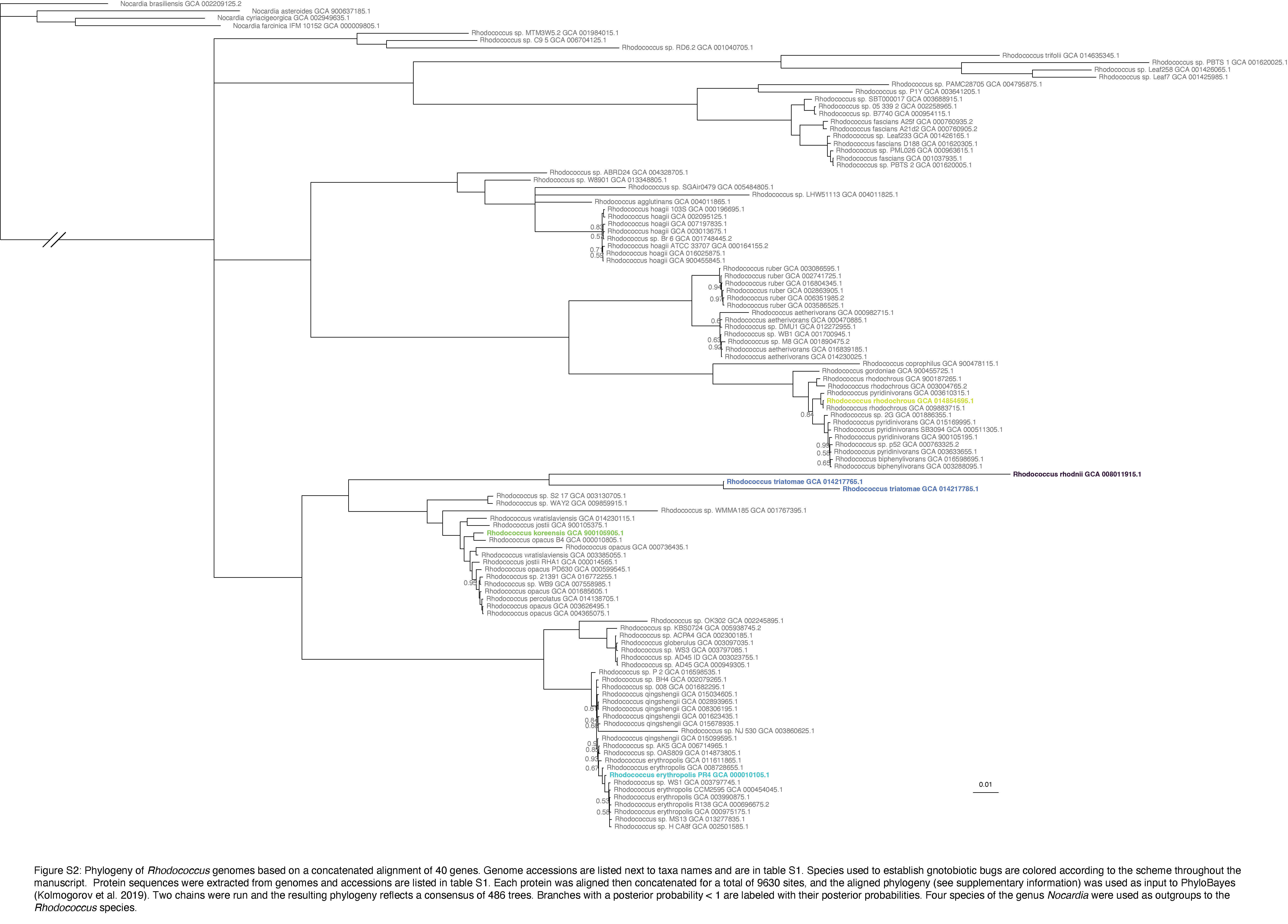
